# Small Molecule-Based Blockade of CD28 Suppresses T Cell Costimulation Across Cellular and Mucosal Co-culture Models

**DOI:** 10.1101/2025.07.16.665260

**Authors:** Saurabh Upadhyay, Valerij Talagayev, Sungwoo Cho, Gerhard Wolber, Moustafa Gabr

## Abstract

**Background and Purpose:** CD28 is a pivotal costimulatory receptor that governs T cell activation through interaction with B7 ligands (CD80/CD86). While antibody-based inhibitors of CD28 signaling have advanced clinically, the development of small molecule modulators remains limited due to the receptor’s shallow, flexible surface. We sought to discover small-molecule modulators with favorable pharmacokinetic properties capable of disrupting CD28–B7 interactions in translational models of T cell activation.

**Experimental Approach:** Structure-based virtual screening was conducted using pharmacophore filtering and consensus docking against the human CD28 ectodomain. Hit compounds were validated using temperature-related intensity change (TRIC) and microscale thermophoresis (MST). Functional antagonism was assessed through ELISA, NanoBit luciferase complementation, and a CD28 Blockade Bioassay. In vitro ADME and safety pharmacology profiling were performed, and immunosuppressive activity was evaluated in tumor–PBMC and mucosal-PBMC co-culture assays.

**Key Results:** Lead compound 22VS bound CD28 in biophysical screening, targeting a lipophilic canyon anchored by K24, Q25, and P27. 22VS inhibited CD28–CD80/CD86 interactions in ELISA and cell-based assays with submicromolar potency. 22VS robustly suppressed T cell activation markers in both tumor– PBMC and human mucosal epithelial–PBMC co-culture models, phenocopying the anti-CD28 biologic FR104. It showed no cytotoxicity up to 300 µM and exhibited high solubility, low clearance, strong membrane permeability, and minimal off-target effects in pharmacokinetic screens.

**Conclusion and Implications:** This study identifies a novel druggable site on CD28 and validates 22VS as a selective, non-toxic small molecule inhibitor with translational potential for immune modulation in autoimmunity, transplantation, and cancer.

**Bullet Point Summary:** *What is already known:*

- CD28 is a key T cell costimulatory receptor essential for immune activation.
- Small-molecule inhibitors of CD28 are largely unexplored compared to biologics.

*What this study adds:*

- Identifies a novel druggable pocket on CD28 via structure-based virtual screening.
- Discovers 22VS, a selective small molecule CD28 inhibitor with cellular activity.
- Demonstrates that 22VS suppresses T cell activation in tumor–PBMC and mucosal-PBMC co-culture assays, phenocopying a benchmark biologic (FR104).
- Establishes 22VS as a drug-like compound with favorable in vitro pharmacokinetic properties, including metabolic stability, permeability, and low off-target toxicity.

*Clinical significance:*

- Highlights the potential of 22VS as a lead for immunomodulatory therapeutic development.
- Supports small-molecule targeting of CD28–B7 interactions in T cell-driven diseases.

**Graphical Abstract:** 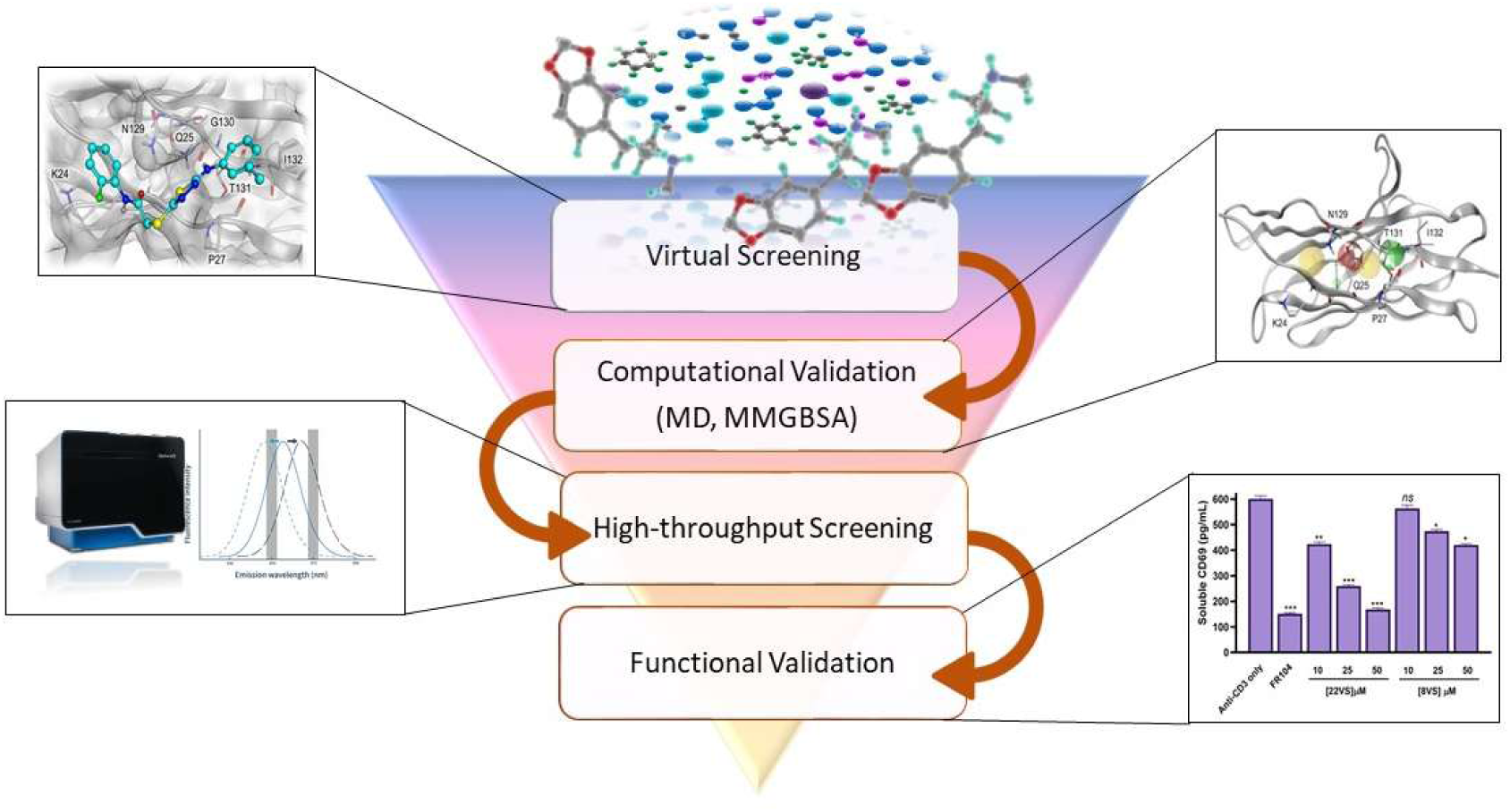

## 1. Introduction

Cluster of Differentiation 28 (CD28) is a key costimulatory receptor expressed on the surface of naïve T cells and is indispensable for full T cell activation, proliferation, survival, and cytokine production (Upadhyay, Kaur, et al., 2025). Upon engagement with its ligands CD80 (B7-1) and CD86 (B7-2) on antigen-presenting cells (APCs), CD28 amplifies T cell receptor (TCR) signaling, leading to robust interleukin-2 (IL-2) secretion and clonal expansion (Levine et al., 1995). While this signaling axis is essential for immune surveillance and pathogen clearance, sustained or unregulated CD28-mediated activation contributes to several pathological conditions, including autoimmunity, graft-versus-host disease, chronic inflammation, and tumor progression (Lai et al., 2021; Linsley & Nadler, 2009; Salomon et al., 2000; Yu et al., 1998). As such, the CD28–B7 interaction has emerged as an attractive immunotherapeutic target in both stimulatory and inhibitory contexts.

Several antibody-based agents have been developed to modulate the CD28 pathway. The most well-known is TGN1412, a superagonist antibody intended to expand regulatory T cells; however, it caused a life-threatening cytokine storm in a Phase I clinical trial, underscoring the risks associated with broad CD28 activation (Suntharalingam et al., 2006). In contrast, antagonistic antibodies such as FR104, a monovalent pegylated Fab’ fragment, and Lulizumab pegol (BMS-931699) have shown favorable immunosuppressive effects in preclinical models of autoimmunity by selectively blocking CD28 without affecting CTLA-4 engagement (Poirier et al., 2015; Shi et al., 2017). Additionally, Abatacept and Belatacept, CTLA-4-Ig fusion proteins that outcompete CD28 for B7 ligands, are already in clinical use for rheumatoid arthritis and transplant rejection (Ostor, 2008), (Linsley et al., 1992). However, these biologics face limitations such as immunogenicity, high production costs, parenteral administration, limited tissue penetration, and systemic immunosuppression (Chames et al., 2009; Feng et al., 2024).

Despite its therapeutic potential, CD28 has historically been viewed as undruggable by small molecules due to its flat, solvent-exposed interface and the lack of a well-defined binding pocket (Calvo-Barreiro & Gabr, 2025). Nonetheless, recent advances in structure-guided drug design and computational modeling have made it possible to explore transient or cryptic pockets within protein–protein interaction interfaces (Calvo-Barreiro et al., 2025). Small molecule inhibitors offer several key advantages over antibody-based therapies, including oral bioavailability, lower production costs, superior tissue and tumor penetration, reduced immunogenicity risk, and reversible pharmacodynamics. These properties render small molecules uniquely suited for chronic conditions, solid tumors, and therapeutic applications requiring broad tissue access—clinical contexts where biologics may fall short.

In this study, we undertook a structure-guided ligand discovery campaign to identify small-molecule inhibitors of CD28. Starting with the highest-resolution crystal structure of CD28 in complex with a mitogenic antibody Fab fragment (Evans et al., 2005), we performed molecular dynamics (MD) simulations and Pyrod- based water network analysis to uncover a previously unrecognized lipophilic canyon at the CD28 interface (Schaller et al., 2019). A pharmacophore model derived from this site was used to screen over 7 million compounds from the Enamine collection (Wolber & Langer, 2005). Among the validated ligands, compound 22VS demonstrated the most promising CD28 binding affinity and submicromolar inhibition of CD28–B7 interactions in ELISA, NanoBit complementation, and a luciferase-based Jurkat-Raji co-culture assay, with minimal cytotoxicity.

This work highlights the feasibility of drugging a traditionally intractable immune receptor and establishes a validated chemical starting point for the development of selective CD28–B7 inhibitors. Notably, this strategy aligns with a broader trend in drug discovery where challenging protein–protein interfaces and atypical binding sites—such as those in PKM2, G protein–coupled receptors (GPCRs), and D-amino acid oxidase (DAAO)—are being successfully exploited using structure-guided design and dynamic binding site mapping (Khan et al., 2024; Marino & Filizola, 2018; Upadhyay, Bhardwaj, et al., 2025). As interest grows in fine-tuning immune responses through PPI blockade, our findings offer timely and translationally relevant insights into next generation immunomodulators.

## 2. Methods

### 2.1 Structure Preparation and Molecular Dynamics Simulations

The structure preparation started with the crystal structure with PDB ID: 1YJD (Evans et al., 2005) . Structure preparation was performed with MOE 2022.02 (Chemical Computing Group, Montreal, Canada). The structure was prepared after the removal of the co-crystalized antibody. The crystallographic water and 2-acetamido-2-deoxy-beta-D-glucopyranose (NAG) covalently bound to CD28 were removed. The structure was protonated at pH 7 and temperature of 300 K using Protonate 3D function implemented in MOE 2022.02 (Labute, 2009).

CD28 was prepared for molecular dynamics (MD) simulations with the use of Maestro 11.7 (Schrödinger, LLC, New York, USA). The hydrogen bond network was optimized at pH 7.0. The protein was placed in an orthorhombic box keeping the edges in a 10 Å distance to the protein surface. The box was filled with TIP3P (Mark & Nilsson, 2001) water model, sodium, and chloride ions to neutralize the system and achieve isotonic conditions (0.15 M NaCl) (Harrach & Drossel, 2014). The MD simulations were performed in 10 replicates, 10 ns each, generating 1000 frames per replica. The system was simulated as NPT ensembles keeping the number of particles, pressure, and temperature constant (p= 1.01325 bar, T= 298 K). The temperature was controlled using the Nose-Hoover thermostat and the pressure using the Martyna-Tobias-Klein method (DeVane et al., 2003; Kleinerman et al., 2008). The MD simulations were carried out with Desmond in version 2022-1 on RTX2080Ti and 3090 graphics processing units (NVIDIA Corporation, Santa Clara, USA).

Pyrod (Schaller et al., 2019) analysis was performed with the grid parameters being selected as the 0, 0, 0 coordinates with the edge lengths being: 45, 50, 60. The pharmacophore feature selection relied on the identification of hydrophobic features. For the identification of the appropriate pharmacophores a threshold was set on hydrogen bond acceptors of 12.7 kcal/mol, hydrogen bond donors 9.17 kcal/mol and the hydrophobic features of 8.3 kcal/mol.

### 2.2 Pharmacophore-based Virtual Screening

The 3D pharmacophore was used for a virtual screening campaign of the Enamine Screening collection database (version 2024, retrieved from enamine.com) consisting of 7.1 million compounds. The library was prepared using idbgen implemented in Ligandscout 4.4.3 with the settings generating 25 conformations for each ligand with a minimal surface accessibility threshold for hydrophobicity score of 0.25 (Wolber & Langer, 2005). Virtual screening was performed using iscreen of Ligandscout 4.4.3 with the default settings that consist of a minimum number of required features of 3 with 0 allowed features to omit. The scoring function used in iscreen was the absolute scoring function.

### 2.3 Molecular Docking Studies

The virtual screening hits underwent molecular docking studies with GOLD 5.8.2 (Jones et al., 1997). The binding site was defined as a sphere of 12 Å radius with a center on the CA atom of T131. The workflow consisted of two molecular docking runs. During the first run, ten genetic algorithm runs per molecule were conducted using the ChemPLP scoring function (Korb et al., 2009). The interaction features of the obtained docking poses were calculated and molecules contained at least two hydrophobic contacts, one hydrogen bond acceptor and one hydrogen bond donor interaction were retained. Those docking poses were then filtered in accordance with their fulfillment of the 3D pharmacophore used during the virtual screening. Next the molecules were minimized with the MMFF94 force field and were again filtered according to their interaction features and 3D pharmacophore fulfillment (Halgren, 1999). This resulted in 778 molecules, which underwent the second round of molecular docking studies. During this stage, ten genetic algorithm runs per molecule were conducted using the Chemscore scoring function (Verdonk et al., 2003). The docking poses were filtered identically as in the previous step with according to their interaction features and fulfillment of the 3D pharmacophore. The final step consisted in the visual inspection of the docking poses with an emphasis on the protein-ligand interaction and the conformational and structural sanity of the obtained docking poses, resulting in 33 unique molecules.

### 2.4 Screening of Small-Molecule Binders to Human CD28 Using Dianthus TRIC Assay

Human CD28 protein (His-tagged; Acro Biosystems) was labeled with RED-tris-NTA 2nd Generation dye (NanoTemper Technologies, Cat #MO-L018) according to the manufacturer’s instructions. A solution containing 20 nM dye and 40 nM CD28 was prepared in assay buffer (PBS with 0.005% Tween-20, pH 7.4) and incubated at room temperature for 30 minutes in the dark.

For high-throughput screening, the labeled CD28 was mixed in a 1:1 ratio with test compounds (final concentration 100 µM in 2% DMSO) or control solutions and incubated for 15 minutes at room temperature in the dark. Human CD80 (2 µM) was used as the positive control, while 2% DMSO alone served as the negative control. After brief centrifugation (1 minutes for 1000g), 20 µL samples were loaded into the Dianthus NT.23 Pico (NanoTemper Technologies) and analyzed at 25 °C. Fluorescence at 670 nm was recorded for 1 second before and 5 seconds after IR laser activation. The normalized TRIC signal (Fnorm) was calculated as the ratio of post-laser (Fhot) to pre-laser (Fcold) fluorescence. In the single-dose experiments, each compound was tested in three technical replicates, while the positive and negative controls were included in duplicate at regular intervals across the 384-well plate.

### 2.5 Monolith: Binding Affinity Screening Platform

As in the HTS using the Dianthus NT.23 Pico (NanoTemper Technologies), the Monolith His-Tag Labeling Kit RED-tris-NTA 2nd Generation was used to label His-tagged human CD28 protein (Acro Biosystems), following the manufacturer’s instructions. A solution of 50 nM RED-tris-NTA dye and 100 nM CD28 protein was prepared in assay buffer (1× PBS with 0.005% Tween-20, pH 7.4) and incubated at room temperature for 30 minutes in the dark before each experiment.

For binding affinity experiments, 12-point serial dilutions of each candidate compound (starting at 500 µM) were prepared in assay buffer with 2% DMSO to yield 2× concentrations. Labeled protein and compound were mixed at a 1:1 ratio (final volume 15 µL), incubated for 15 minutes at room temperature in the dark, and homogenized by pipetting. Samples were then loaded into Monolith Capillaries (Cat# MO-K022) and analyzed using the Monolith NT.115 (NanoTemper Technologies) at 25 °C, with 40% excitation and medium MST power.

Fnorm was calculated as the ratio of fluorescence after to before IR laser activation. Each compound was tested in four technical replicates. For confirmed binders, binding affinities were determined in three independent experiments. Data were analyzed using MO.Affinity Analysis and GraphPad Prism 10.

### 2.6 Evaluation of CD28–CD80 Binding Inhibition via Competitive ELISA

To evaluate the ability of small molecules to disrupt the CD28–CD80 interaction, a competitive ELISA was performed using the CD28:B7-1 [Biotinylated] Inhibitor Screening Assay Kit (Cat# 72007; BPS Bioscience, San Diego, CA, USA), following the manufacturer’s instructions.

Briefly, 96-well plates were coated with CD28 (2 µg/mL in PBS, 50 µL/well) and incubated overnight at 4 °C. The next day, wells were washed with 1× Immuno Buffer and blocked with Blocking Buffer for 1 hour at room temperature. Test compounds at varying concentrations were co-incubated with biotinylated CD80 (B7-1, 5 ng/µL) for 1 hour to allow competitive binding to immobilized CD28. Wells without coating served as ligand controls, while wells treated with inhibitor buffer instead of compound served as negative controls.

Following incubation, plates were washed and incubated with Streptavidin-HRP (1:1000 dilution in Blocking Buffer) for 1 hour. After a final wash, chemiluminescent substrate was added, and luminescence was measured immediately using a plate reader in luminescence mode.

### 2.7 Cell viability assay

The cytotoxic effects of test compounds were evaluated using the MTS assay (Cell Titer 96® AQueous One Solution, Promega, WI, USA). Jurkat human T lymphocyte cells were seeded in 96-well plates and maintained in complete growth medium containing 10% FBS until reaching ap-proximately 40% confluence (24 h). Cells were then treated with compounds and Tween-20 for 24 h followed by MTS analysis according to the supplier’s protocol. Absorbance readings were obtained at 490 nm using an Infinite M1000 Pro Microplate Reader (Tecan, Männedorf, Switzerland).

### 2.8 NanoBit assay

For the measurement of protein-protein interactions, CHO-K1 cells were transfected with plasmids encoding CD28 fused to a C-terminal SmBiT and CD80 or CD86 fused to a C-terminal LgBiT(van der Merwe et al., 1997). Transfected cells were seeded in 96-well white microplates and cultured until reaching approximately 70% confluence. Prior to drug treatment, cells were washed twice with phosphate-buffered saline (PBS) to remove residual culture medium. Each well was then filled with 50 μL of Hank’s Buffered Salt Solution (HBSS), and test compounds were added to the designated wells. Cells were incubated with the respective drugs for 2 hours at 37°C. Following drug treatment, furimazine substrate was added to each well at a final concentration of 10 μM and incubated for 10 minutes at room temperature. Luminescence signals were measured using an Infinite M1000 Pro Microplate Reader (Tecan, Männedorf, Switzerland). Background luminescence was subtracted from all measurements, and relative luminescence units (RLU) were calculated to determine the extent of protein-protein interactions. All experiments were performed in triplicate, and data represent the mean ± standard error of the mean (SEM) from at least three independent experiments.

### 2.9 CD28 Blockade Bioassay in 96-Well Format with Small-Molecule Screening and Human Serum Tolerance

The CD28 Blockade Bioassay (Promega) was adapted to a 96-well format to screen small molecules for their ability to inhibit CD28-mediated costimulation. Test compounds were prepared in a 10-point, 1:1 serial dilution starting at 200 µM in assay buffer containing 1% DMSO. Each dilution (50 µL/well) was added to white 96-well plates.

CD28 Effector Cells (2 × 10⁴ cells/well) were first seeded, followed by preincubation with diluted compound and anti-CD28 control antibody (Cat# K1231, Promega) for 5 minutes at room temperature. Next, aAPC/Raji Cells (2 × 10⁴ cells/well) were added, and plates were incubated at 37 °C with 5% CO₂ for 5 hours. After incubation, 50 µL of Bio-Glo™ Reagent (Promega) was added to each well, and luminescence was recorded using a GloMax® Discover System.

Dose–response curves were generated and fitted using a four-parameter logistic regression in GraphPad Prism 10 to calculate percent inhibition and IC_50_ values. The assay showed robust performance in the presence of up to 10% pooled human serum, indicating tolerance to physiological matrix conditions.

### 2.10 Pharmacokinetic profiling

The preliminary evaluation of PK parameters for compounds 22VS and 8VS was performed as previously reported by us (Kaur et al., 2025).

### 2.11 Evaluation of T Cell Activation in Tumor-PBMC Coculture

To assess the dose-dependent impact of CD28 inhibition on T cell activation in a translational human system, we employed a 3D tumor–PBMC co-culture assay. Tumor spheroids were generated by seeding A549 (or MDA-MB-231) cells in ultra-low attachment 96-well plates. After 48 hours, spheroids were treated with recombinant human interferon-gamma (IFN-γ, 50 ng/mL) for 24 hours. Freshly thawed human peripheral blood mononuclear cells (PBMCs) were added at a 5:1 effector-to-target (E:T) ratio, and co-cultures were stimulated with soluble anti-CD3 antibody (0.3 μg/mL, clone OKT3) to model TCR activation.

Compounds were added at the initiation of co-culture. FR104 (10 μg/mL), a reference anti-CD28 Fab’ biologic, served as a positive control. The test compounds 22VS and 8VS were evaluated at three concentrations (10, 25, and 50 μM). Vehicle and anti-CD3–only conditions were included as controls. After 48 hours, supernatants were harvested and analyzed for IFN-γ and IL-2 levels using human-specific ELISA kits (BioLegend). Soluble CD69 (sCD69), released from activated T cells, was quantified using a human CD69 ELISA kit (ThermoFisher).

### 2.12 CD28-Dependent Cytokine Release in a Human PBMC–Mucosal Co-culture Model

To evaluate the immunomodulatory activity of 22VS in a mucosal-relevant human immune interface, we employed a commercially available human airway epithelial tissue model (MucilAir™, Epithelix) co-cultured with PBMCs. Cryopreserved human PBMCs (StemCell Technologies) were thawed and rested for 4 hours in RPMI-1640 medium supplemented with 10% heat-inactivated human AB serum, 2 mM L-glutamine, and 1% penicillin–streptomycin.

MucilAir™ inserts were equilibrated in the manufacturer’s proprietary maintenance medium for 24 hours at 37 °C and 5% CO2. PBMCs were resuspended at 2 × 10^6^ cells/mL and overlaid (200 µL per insert) onto the apical surface of the MucilAir™ tissue in 24-well Transwell plates. Co-cultures were stimulated with plate-bound anti-CD3 (1 µg/mL) and soluble anti-CD28 (1 µg/mL) in the presence of: vehicle control (0.1% DMSO), 22VS (10, 25, or 50 µM), or FR104 (10 µg/mL; anti-CD28 Fab’ fragment).

After 48 hours, apical supernatants were harvested and analyzed for IFN-γ, IL-2, and TNF-α concentrations using human ELISA kits (BioLegend) according to manufacturer instructions. Cytokine concentrations were interpolated from standard curves and expressed in pg/mL. All conditions were performed in triplicate (three independent experiments).

## 3. Results

### 3.1 Structure-Based Identification and Hit Discovery for CD28

To identify novel small molecule binders of human CD28, a structure-based virtual screening (VS) campaign was initiated using the crystal structure of CD28 in complex with a mitogenic Fab antibody (PDB ID: 1YJD; 2.7 Å resolution (Evans et al., 2005)) (Figure 1A). After protein preparation and molecular dynamics (MD) simulations in explicit solvent, Pyrod (Schaller et al., 2019) was applied to analyze dynamic water-mediated interactions. This analysis revealed a previously uncharacterized lipophilic canyon on the CD28 surface, enriched in druggable interaction hotspots, including two hydrophobic features, two hydrogen bond donors, and one acceptor, located around residues L22, K24, Q25, P27, I109, N129, G130, T131, and I132 (Figure 1B-D).

**Figure 1.**
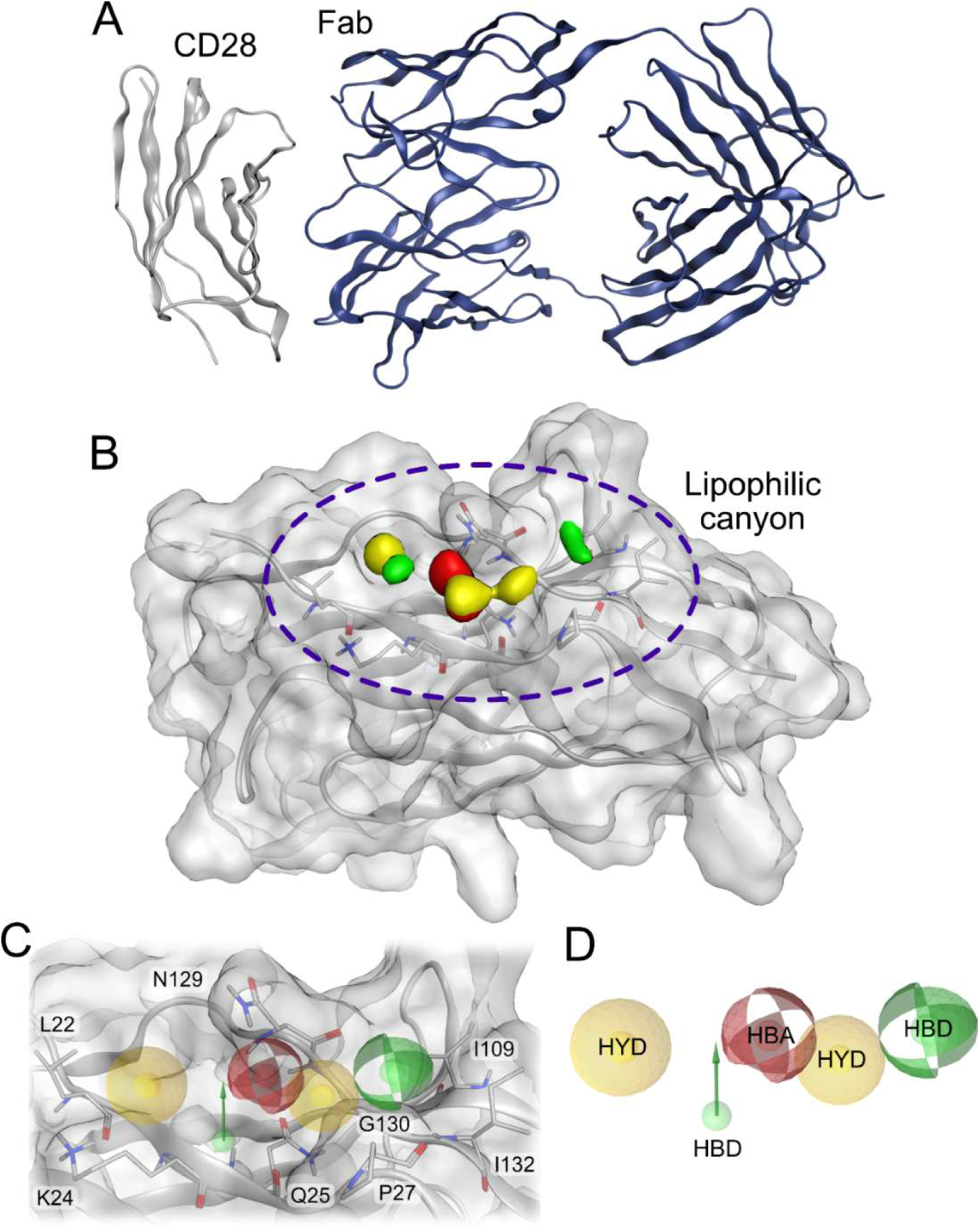
Structure-based identification of a lipophilic binding site on CD28 and virtual screening results. **(A)** The crystal structure of human CD28 (grey ribbon) in complex with the Fab fragment of a mitogenic antibody (dark blue ribbons), from PDB ID: 1YJD (Evans et al., 2005), served as the structural basis for virtual screening. **(B)** Pyrod analysis (Schaller et al., 2019) of MD simulations revealed a distinct lipophilic canyon on CD28, visualized using dynamic molecular interaction fields (dMIFs), where yellow spheres represent hydrophobic features, red spheres indicate hydrogen bond acceptors, and green spheres indicate hydrogen bond donors. **(C)** A 3D pharmacophore was generated from these features; the upper panel shows a close-up of the pharmacophoric features within the lipophilic canyon, with the hydrogen bond acceptor being located next to N129 and the hydrogen bond donors are in close proximity to Q25 and G130. One hydrophobic feature is next to the side chain of L22, while the second hydrophobic feature is next to I109. **(D)** the generated pharmacophore model consisting out of two hydrophobic features, two hydrogen bond donors and one hydrogen bond acceptor was used to screen the Enamine compound library.

A 3D pharmacophore was constructed based on these features and used to screen the Enamine Screening Collection (∼7.15 million compounds) using LigandScout 4.4.3 (Gerhard Wolber & Thierry Langer, 2005). The screen yielded 32,097 initial hits. After physicochemical filtering and a two-step consensus docking protocol using both GOLD (Gareth Jones et al., 1997), 33 top-ranking compounds were selected for further validation. A recurring 1,3,4-thiadiazole scaffold was observed in a subset of hits, suggesting a privileged chemotype for interaction with the CD28 pocket.

### 3.2 Biophysical Screening and Affinity Profiling Using TRIC and MST

To experimentally validate binding, the 33 compounds were subjected to a two-stage screening protocol. In the first stage, TRIC-based high-throughput screening was performed on the Dianthus NT.23 Pico instrument. Labeled CD28 protein was incubated with each compound at 100 µM in assay buffer containing 4% DMSO. Each compound was tested in three technical replicates, and 32 negative controls (buffer + DMSO) were evenly distributed across the plate to monitor laser drift and background variability.

Normalized fluorescence (Fnorm) values were calculated for each compound. Five compounds exceeded the ±5 SD threshold relative to the negative control mean (Figure 2A), indicating potential binding events. These five candidates were further evaluated by MST for dose-dependent binding.

**Figure 2.**
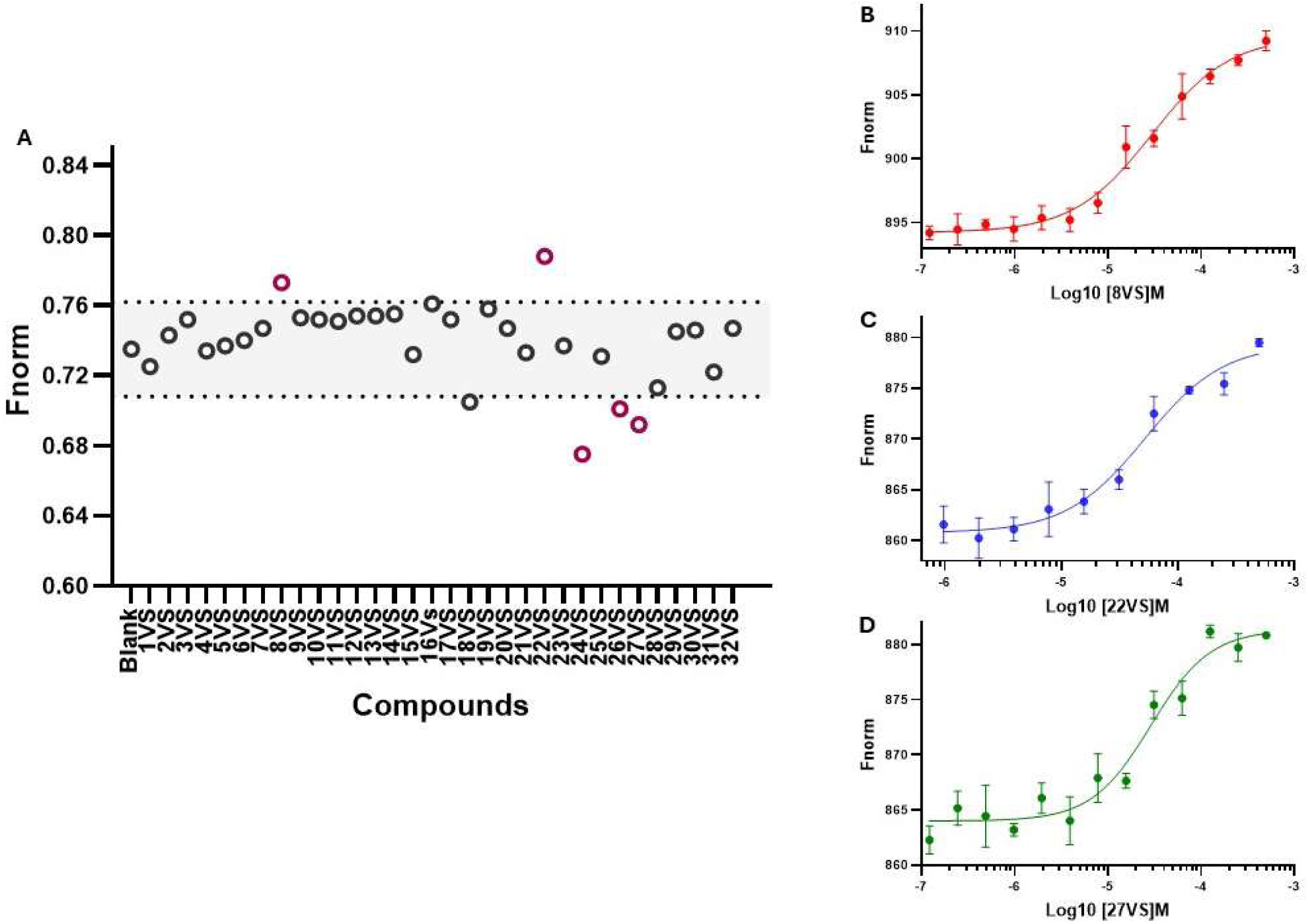
Biophysical identification and validation of CD28-binding small molecules using TRIC-based Dianthus screening and microscale thermophoresis (MST). **(A)** Single-dose TRIC-based screening of small-molecule compounds against CD28 protein using the Dianthus platform. Each compound was tested at 100 μM in the presence of 4% DMSO, with CD28 alone serving as the negative control. The black dotted lines represent a ±5-fold standard deviation threshold relative to the negative control. Compounds falling within this range were considered non-binders, while those outsides were classified as potential hits. (b–d) MST-based dose–response validation of three compounds (8VS, 22VS, and 27VS) that showed positive signals in the Dianthus screen. MST measurements were performed using the same buffer conditions with 4% DMSO. Sigmoidal binding curves confirmed specific and concentration-dependent interactions with CD28, yielding dissociation constants (Kd) of 29.57 ± 12.69 μM for 8VS **(B)**, 52.45 ± 18.69 μM for 22VS **(C)**, and 38.35 ± 17.14 μM for 27VS **(D)**. The resulting data were fitted using a nonlinear regression model—log(inhibitor) vs. response with a variable slope (four parameters)—using GraphPad Prism software. Error bars represent the standard deviation from three replicates.

Microscale thermophoresis (MST) confirmed specific binding for three compounds: 8VS, 22VS, and 27VS. Each compound was tested in a 16-point serial dilution (starting at 500 µM), and MST responses were recorded in three independent experiments. Sigmoidal binding curves were obtained with dissociation constants (Kd) of 29.57 ± 12.69 µM for 8VS, 52.45 ± 18.69 µM for 22VS, and 38.35 ± 17.14 µM for 27VS (Figure 2B–D). These values indicated moderate micromolar affinity. The other two compounds failed to show reproducible, dose-dependent binding, suggesting non-specific effects or weak interactions. Chemical structures of the identified hits are displayed in Figure 3.

**Figure 3.**
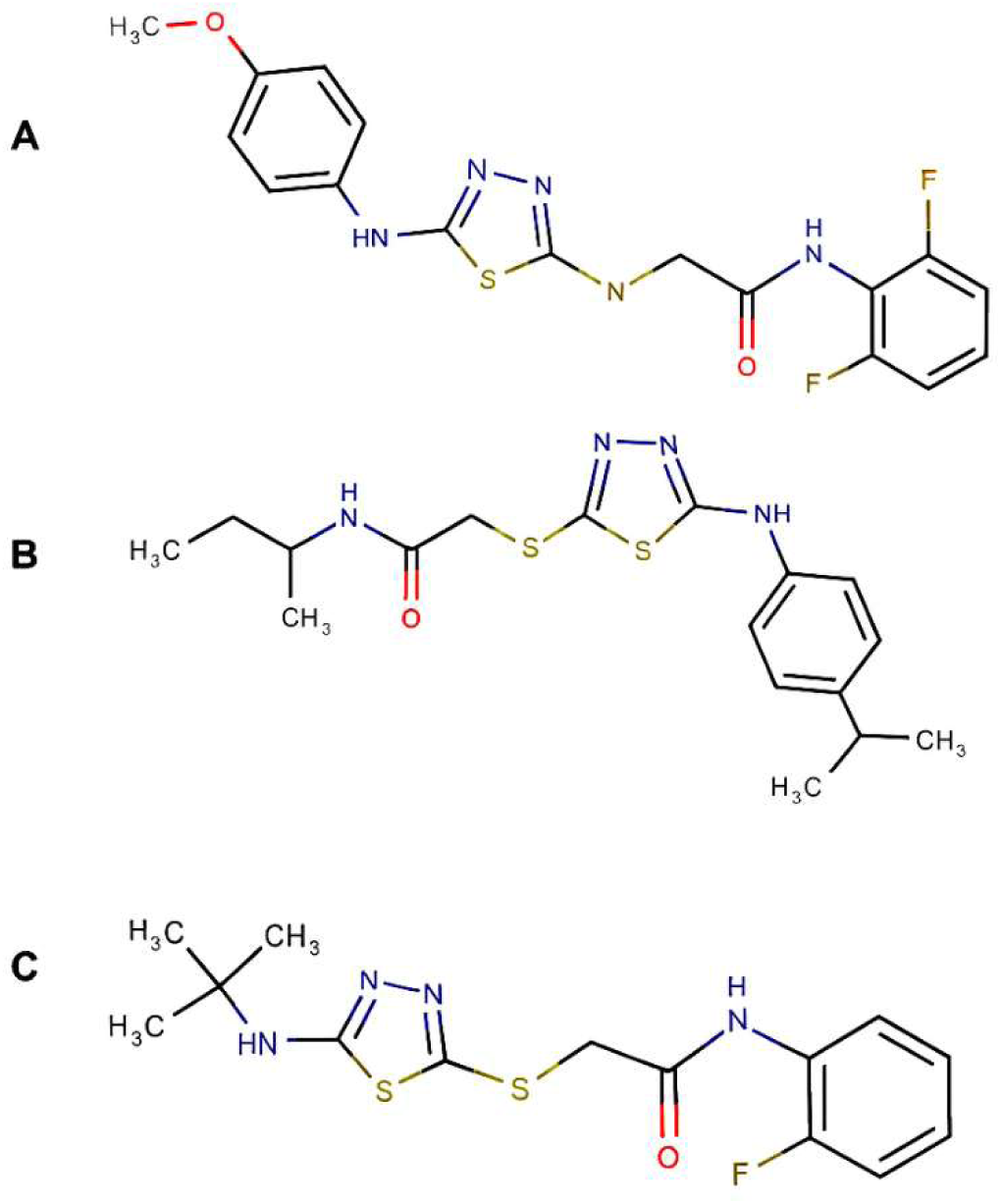
Chemical structures of CD28-targeting small molecules identified through structure-guided screening. **(A)** 8VS, **(B)** 22VS, and **(C)** 27VS represent distinct scaffolds incorporating triazole- thioamide linkages and variable terminal groups designed to engage a lipophilic pocket on CD28 and disrupt B7-mediated co-stimulatory signaling.

### 3.3 Functional Inhibition of CD28–CD80 Interaction in ELISA

To assess the ability of biophysically validated compounds to functionally disrupt CD28-ligand binding, an ELISA-based CD28:B7-1 interaction assay was conducted. Compounds 8VS and 22VS were selected based on solubility and MST profiles. CD28 was immobilized on 96-well plates and incubated with biotinylated CD80 in the presence of increasing compound concentrations. Both compounds inhibited CD80 binding in a concentration-dependent manner.

Compound 22VS showed potent inhibition with an IC50 of 7.80 ± 2.47 µM, while 8VS exhibited moderate inhibition with an IC50 of 65.45 ± 20.89 µM (Figure 4A–B). These values aligned well with MST-derived affinities and provided initial evidence for functional blockade of the CD28:B7-1 axis.

**Figure 4.**
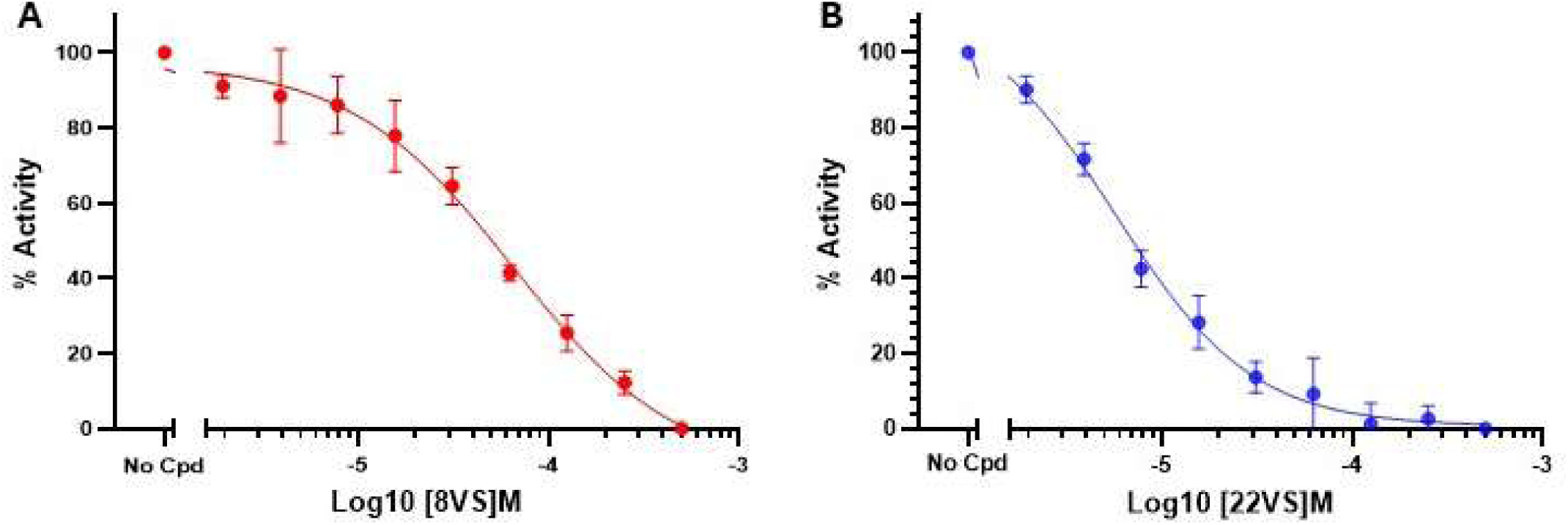
**Inhibition of CD28:B7-1 interaction by virtual screening-derived small molecules using the ELISA-based inhibitor screening assay**. **(A)** Dose-response curve of compound 8VS showing an IC₅₀ of 65.45 ± 20.89 μM, indicating moderate inhibition of CD28:B7-1 binding. **(B)** Dose-response curve of compound 22VS displaying a significantly lower IC₅₀ of 7.80 ± 2.47 μM, suggesting higher potency in blocking the CD28:B7-1 interaction. The assay was conducted using a 96-well plate format with immobilized CD28 protein and biotinylated B7-1, followed by detection with streptavidin-HRP and chemiluminescence measurement.The resulting data were fitted using a nonlinear regression model—log(inhibitor) vs. response with a variable slope (four parameters)—using GraphPad Prism software.

### 3.4 In silico analysis of compound binding hypothesis

Visual inspection of the binding hypothesis for compounds 8VS and 22VS revealed conserved interaction patterns. Both engage in hydrophobic contacts with the side chains of L22 and K24 on the left side of the lipophilic canyon, as well as with the side chain of I132 on the right side. Additionally, 8VS establishes a hydrophobic contact with I109, while 22VS shows a hydrophobic contact with P27. A hydrogen bond forms between the side chain of N129 and a nitrogen atom of the 1,3,4-thiadiazole scaffold in both compounds. Both 8VS and 22VS form hydrogen bonds with the backbone carbonyl oxygen of G130. Notably, 8VS establishes an additional hydrogen bond with the backbone amide of Q25 (Figure 5)

**Figure 5.**
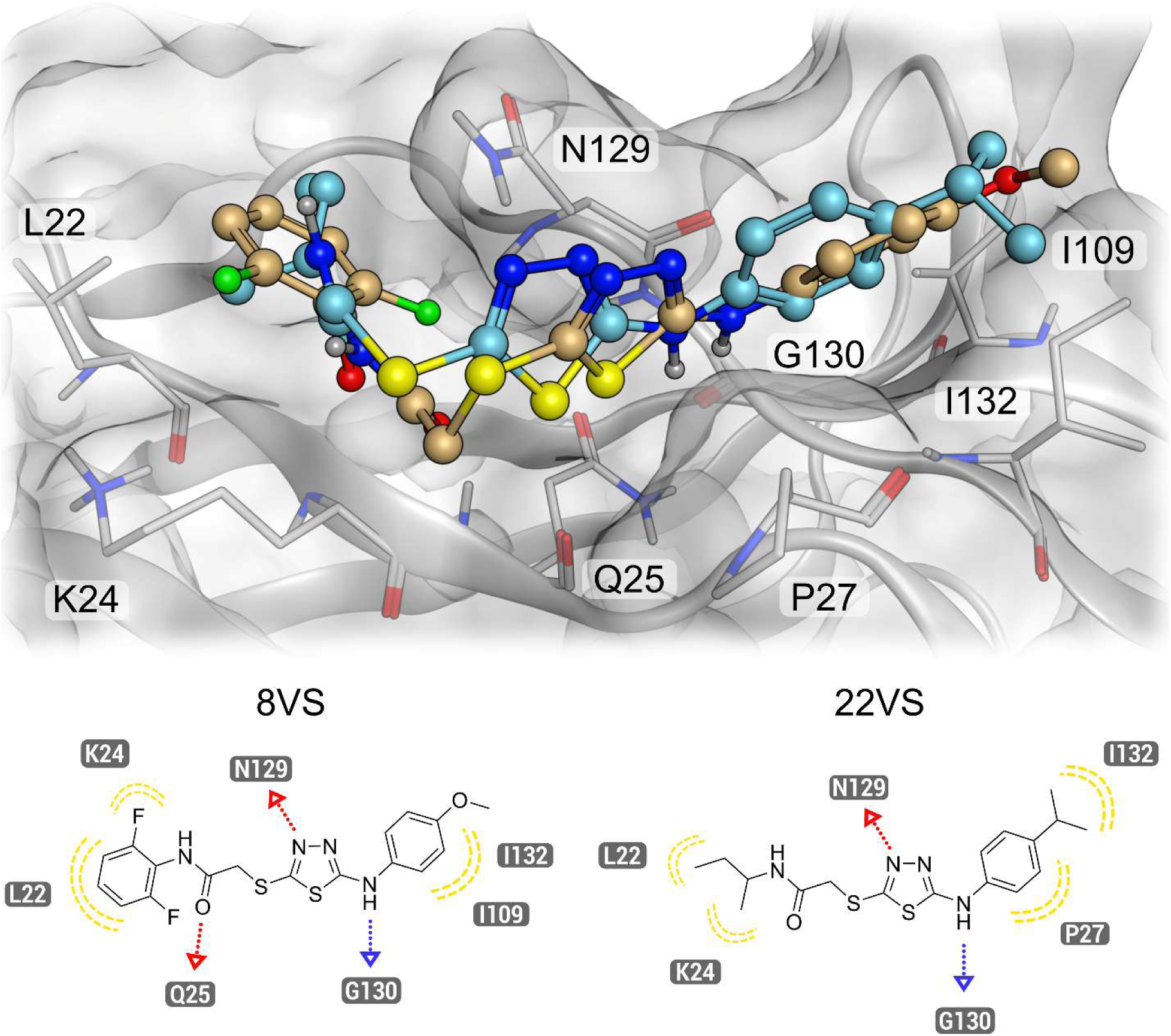
**Binding hypothesis of 8VS and 22VS**. The binding hypothesis of both compounds 8VS and 22VS show a hydrogen bond interaction between the 1,3,4-thiadiazole scaffold and the side chain of N129. 8VS and 22VS show hydrophobic contacts with L22, K24 and I132 with 8VS showing an additional hydrophobic contact with I109 and 22VS with P27. Additionally, both compounds establish a hydrogen bond interaction with G130, while 8VS additionally shows a hydrogen bond interaction with Q25.

### 3.5 Cellular Inhibition of CD28–B7 Interactions and Cytotoxicity Evaluation

To evaluate the ability of small molecules to inhibit CD28–B7 interactions in a cellular context, we employed a NanoBit luciferase complementation assay using CHO-K1 cells co-expressing CD28-SmBiT with either CD80-LgBiT or CD86-LgBiT. Compounds 8VS and 22VS, which showed consistent biophysical binding and ELISA-based functional inhibition, were tested at concentrations up to 100 µM.

Compound 8VS demonstrated minimal inhibitory activity, with no significant suppression of luminescence even at the highest soluble dose tested (Figure 6A–B). In contrast, compound 22VS effectively inhibited both CD28–CD80 and CD28–CD86 interactions in a dose-dependent manner, achieving IC₅₀ values of 1.48 ± 0.3 µM and 1.39 ± 0.4 µM, respectively (Figure 6C–D). The maximal inhibition achieved was ∼30% for CD28–CD80 and ∼40% for CD28–CD86. The slightly enhanced inhibition of CD86 over CD80 may reflect the known differential binding affinities of CD28 for these ligands (CD80: Kd ≈ 4 µM; CD86: Kd ≈ 20 µM), suggesting a ligand-specific modulation profile.

**Figure 6.**
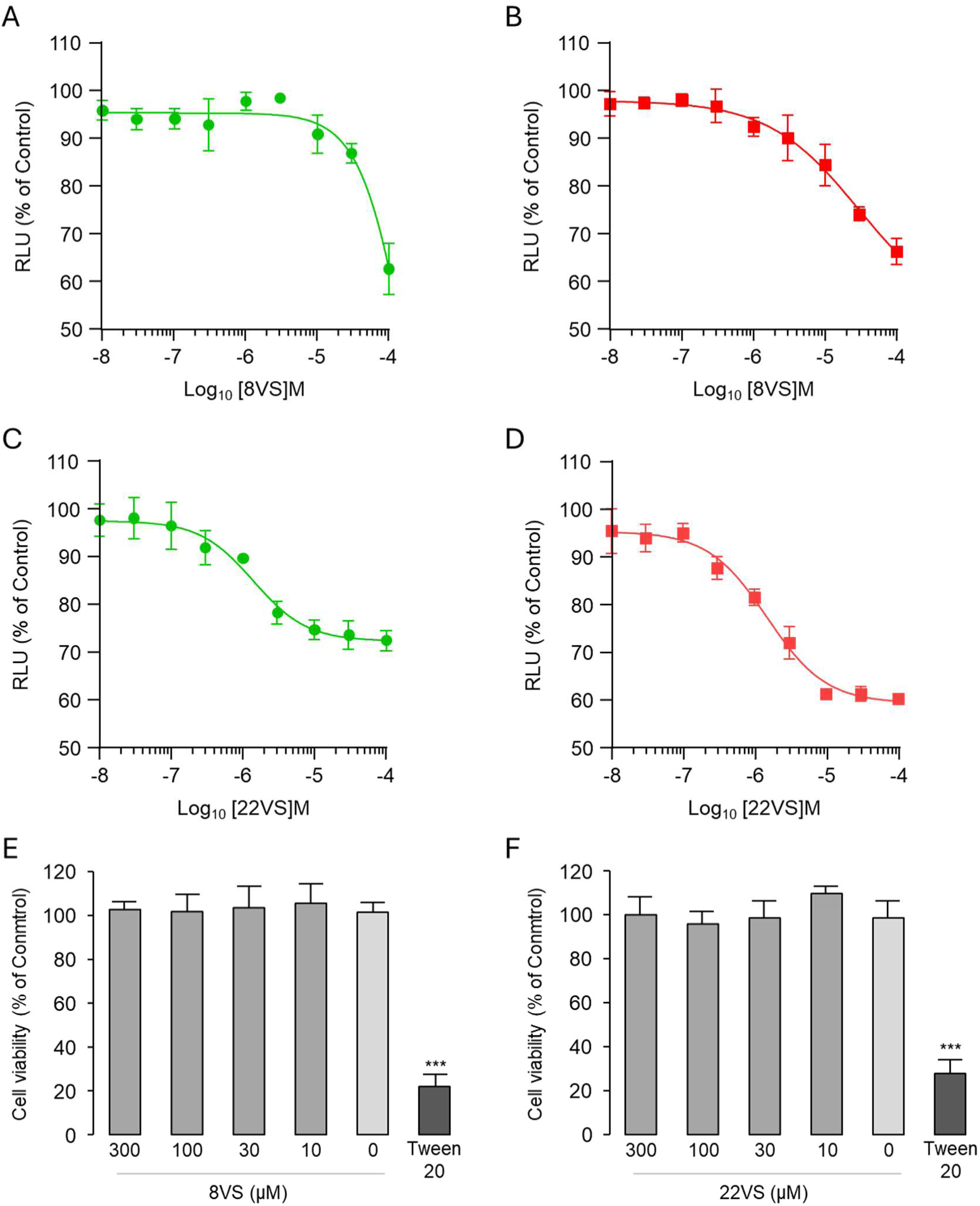
Dose-response analysis and cytotoxicity assessment of 8VS and 22VS compounds. (A, B) Concentration-dependent effects of 8VS on CD28-CD80 **(A)** and CD28-CD86 **(B)** interactions. CHO-K1 cells co-expressing CD28-SmBiT and CD80-LgBiT or CD86-LgBiT were treated with increasing concentrations of 8VS for 2 hours at 37°C. Data show limited inhibitory activity with minimal suppression at 100 μM. (C, D) Dose-response curves for 22VS inhibition of CD28-CD80 **(C)** and CD28-CD86 **(D)** interactions. 22VS demonstrated IC₅₀ values of 1.48 ± 0.3 μM and 1.39 ± 0.4 μM for CD28-CD80 and CD28-CD86 interactions, respectively, with maximum inhibition of 30% and 40%, respectively. **(E, F)** Cell viability assessment in human Jurkat T cells treated with 8VS **(E)** or 22VS **(F)** for 24 hours. Cell viability was measured using MTS assay and expressed as percentage of untreated control. Tween 20 (0.1%) served as positive control for cytotoxicity. Data represent mean ± SEM from three independent experiments. Relative luminescence units (RLU) are expressed as percentage of DMSO-treated control cells. ***p <0.001 vs. untreated control.

To confirm that the observed effects were not due to compound-related cytotoxicity, we performed MTS cell viability assays in human Jurkat T cells exposed to 8VS and 22VS across a concentration range up to 300 µM. Both compounds maintained excellent safety profiles, with cell viability consistently above 85% (Figure 6E–F). In contrast, the positive control (0.1% Tween 20) significantly reduced viability to ∼30%, confirming assay sensitivity. These results support the notion that 22VS specifically inhibits CD28–B7 interactions without inducing nonspecific cytotoxic effects.

### 3.6 Reporter-Based Functional Blockade Assay

The final functional assessment was performed using the CD28 Blockade Bioassay, a luciferase reporter assay in Jurkat Effector Cells co-cultured with Raji aAPC cells. This assay measures the inhibition of CD28- mediated costimulation. Compounds 8VS and 22VS were tested in a dose–response format, and luminescence was measured after 5 hours of incubation.

Both compounds showed clear, dose-dependent inhibition of CD28 signaling (Figure 7A–B). Compound 22VS achieved stronger suppression with an IC50 of 6.96 ± 2.95 µM, while 8VS had an IC50 of 10.84 ± 4.62 µM. These results corroborate prior binding and ELISA data, confirming that both compounds engage CD28 and inhibit downstream signaling.

**Figure 7.**
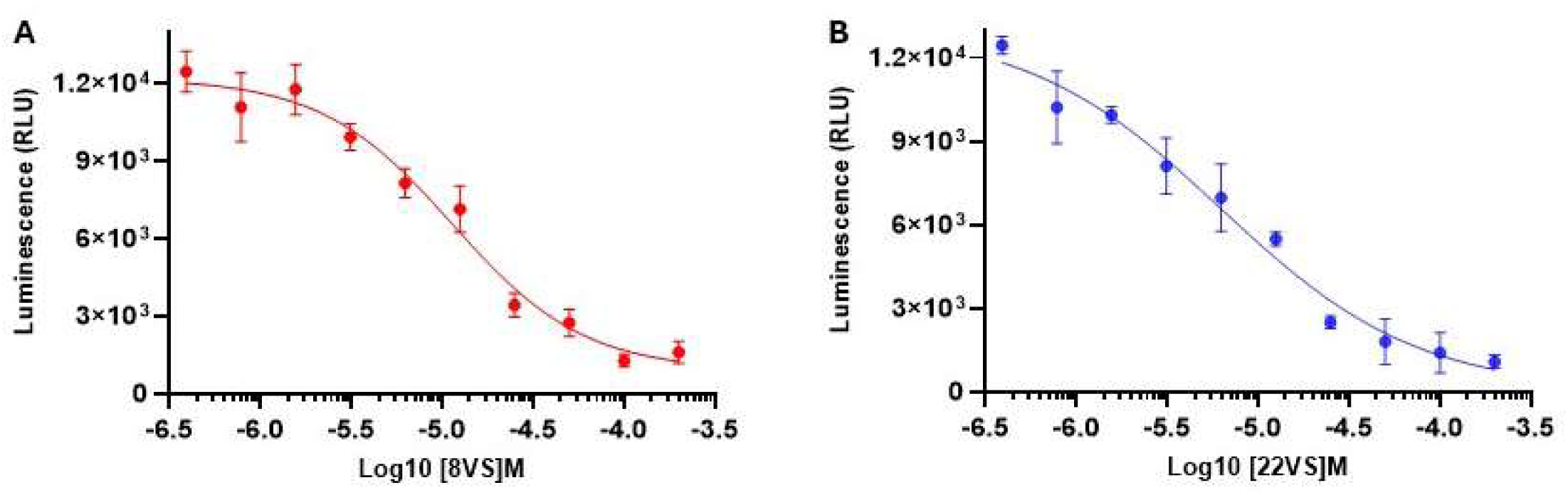
Inhibition of CD28–B7 signaling by small-molecule ligands 8VS and 22VS in a reporter-based blockade assay. Jurkat CD28 Effector Cells and aAPC/Raji Cells were co-cultured in the CD28 Blockade Bioassay (Promega, Cat. #JA6101) to evaluate functional inhibition of CD28–B7 interaction by small molecules. Luminescence was measured using the Bio-Glo™ Luciferase Assay System. **(A)** Compound 8VS inhibited CD28-dependent signaling in a concentration-dependent manner, with an IC₅₀ of 10.84 ± 4.62 µM. **(B)** Compound 22VS showed a stronger inhibitory effect, with an IC50 of 6.96 ± 2.95 µM. Curves were fitted using a four-parameter logistic model in GraphPad Prism, and data are shown as mean ± standard deviation (SD) from n = 3 independent experiments.

### 3.7 Pharmacokinetic (PK) Profiling

We evaluated the in vitro pharmacokinetic (PK) profiles of CD28-targeted compounds 22VS and 8VS to support lead prioritization. As shown in Table 1, 22VS demonstrated superior aqueous solubility—93 µM in PBS and 108 µM in FaSSIF—compared to 8VS (55 µM and 64 µM, respectively). In contrast, 8VS showed higher passive permeability (7.2 × 10⁻⁶ cm/s vs. 4.1 × 10⁻⁶ cm/s), consistent with its greater lipophilicity (LogD₇.₄ = 3.12 vs. 2.95) and lower polarity (TPSA = 66.9 Å² vs. 76.1 Å²). Both compounds were highly stable in simulated gastrointestinal conditions, retaining >80% after 2 hours.

**Table 1.** In vitro PK profiles of compounds 22VS and 8VS.

| PK parameter | 22VS | 8VS |
| --- | --- | --- |
| LogD<sub>7.4</sub><sup>a</sup> | 2.95 | 3.12 |
| TPSA | 66.9 Å<sup>2</sup> | 76.1 Å<sup>2</sup> |
| Kinetic Solubility<br>1% DMSO/PBS<br>[$\mu\text{M}$]<sup>b</sup> | 93 | 55 |
| Solubility in<br>FaSSIF [$\mu\text{M}$] | 108 | 64 |
| PAMPA $P_{app}$<br>[cm/s] | $4.1 \times 10^{-6}$ | $7.2 \times 10^{-6}$ |
| Stability in<br>simulated gastric<br>fluid (pH 1.2, 2h) | >95%<br>remaining | >80%<br>remaining |
| Stability in<br>simulated<br>intestinal fluid<br>(pH 6.8, 2h) | >90%<br>remaining | >85%<br>remaining |
| $t_{1/2}$ [min] mouse<br>liver<br>microsomes<sup>c</sup> /<br>CL<sub>int</sub><sup>d</sup> | $55.1 \pm 4.2$ /<br>$35.7 \pm 2.8$ | $30.8 \pm 1.6$ /<br>$62.4 \pm 5.1$ |
| $t_{1/2}$ [min] human<br>liver<br>microsomes<sup>c</sup> /<br>CL<sub>int</sub><sup>d</sup> | $72.5 \pm 4.1$ /<br>$25.3 \pm 2.3$ | $35.1 \pm 1.3$ /<br>$58.7 \pm 4.2$ |
| $t_{1/2}$ [min] mouse<br>plasma | >100 | $71.3 \pm 5.7$ |
| $t_{1/2}$ [min] human<br>plasma | >100 | $78.1 \pm 3.2$ |
| $t_{1/2}$ [min] PBS | >400 | >250 |
| PPB human [%]<sup>c</sup> | 98.5 ± 0.7 | 94.6 ± 1.1 |
| IC<sub>50</sub> against WI-38<br>[μM] | >100 | >100 |
| IC<sub>50</sub> against HS-27<br>[μM] | >100 | >100 |
| IC<sub>50</sub> against hERG<br>[μM] | >80 | >80 |
| CYP3A4 inhibition (% at 10 mM) | 10.2% | 16.5% |
| CYP2D6 inhibition (% at 10 mM) | 7.4% | 13.2% |
| CYP2C9 inhibition (% at 10 mM) | 6.9% | 12.5% |
| CYP2C19 inhibition (% at 10 mM) | 7.5% | 14.3% |

<sup>a</sup>LogD (pH = 7.4) the distribution coefficient at physiological pH 7.4. <sup>b</sup>Kinetic solubility measured in 1% DMSO/PBS (pH 7.4) by UV-vis spectrophotometry (n=3). <sup>c</sup>Values are mean ± SD (n=3). <sup>d</sup>Intrinsic clearance expressed in μL/(min × mg) protein. Values are mean ± SD (n=3).

Microsomal stability revealed clear differences: in mouse liver microsomes, 22VS showed a t₁/₂ of 55.1 ± 4.2 min and CLint of 35.7 ± 2.8 μL/min/mg, whereas 8VS had a shorter t₁/₂ of 30.8 ± 1.6 min and higher clearance (62.4 ± 5.1 μL/min/mg). Similar trends were observed in human liver microsomes, with 22VS exhibiting better stability (t₁/₂ = 72.5 ± 4.1 min; CLint = 25.3 ± 2.3 μL/min/mg) than 8VS (t₁/₂ = 35.1 ± 1.3 min; CLint = 58.7 ± 4.2 μL/min/mg). Both compounds were stable in plasma and PBS, with 22VS showing plasma half-lives >100 min and buffer stability >400 min; 8VS showed moderate plasma stability (71–78 min) and buffer t₁/₂ >250 min.

Plasma protein binding was high for both (22VS: 98.5 ± 0.7%; 8VS: 94.6 ± 1.1%). Neither compound was cytotoxic in WI-38 or HS-27 fibroblasts (IC₅₀ >100 µM) or inhibited hERG (IC₅₀ >80 µM). However, 22VS showed minimal CYP450 inhibition (6.9%–10.2%), while 8VS exhibited higher inhibition (12.5%–16.5%) across key isoforms.

### 3.8 Evaluation of T Cell Activation in Tumor-PBMC Coculture

We employed a 3D tumor–PBMC co-culture system to assess the functional activity of small-molecule CD28 inhibitors in a physiologically relevant environment. IFN-γ–pretreated A549 tumor spheroids were co- cultured with thawed human PBMCs in the presence of suboptimal anti-CD3 (0.3 μg/mL). Under these conditions, PBMCs exhibited strong activation—marked by elevated levels of IFN-γ (120 pg/mL), IL-2 (100 pg/mL), and soluble CD69 (sCD69, 600 pg/mL)—consistent with CD28 costimulation via tumor-expressed CD80/CD86 (Figures 8A, 8B). The addition of FR104 (10 μg/mL) significantly suppressed activation (IFN-γ = 15 pg/mL, IL-2 = 10 pg/mL, sCD69 = 150 pg/mL), confirming CD28-dependence (Figures 8A, 8B).

**Figure 8.**
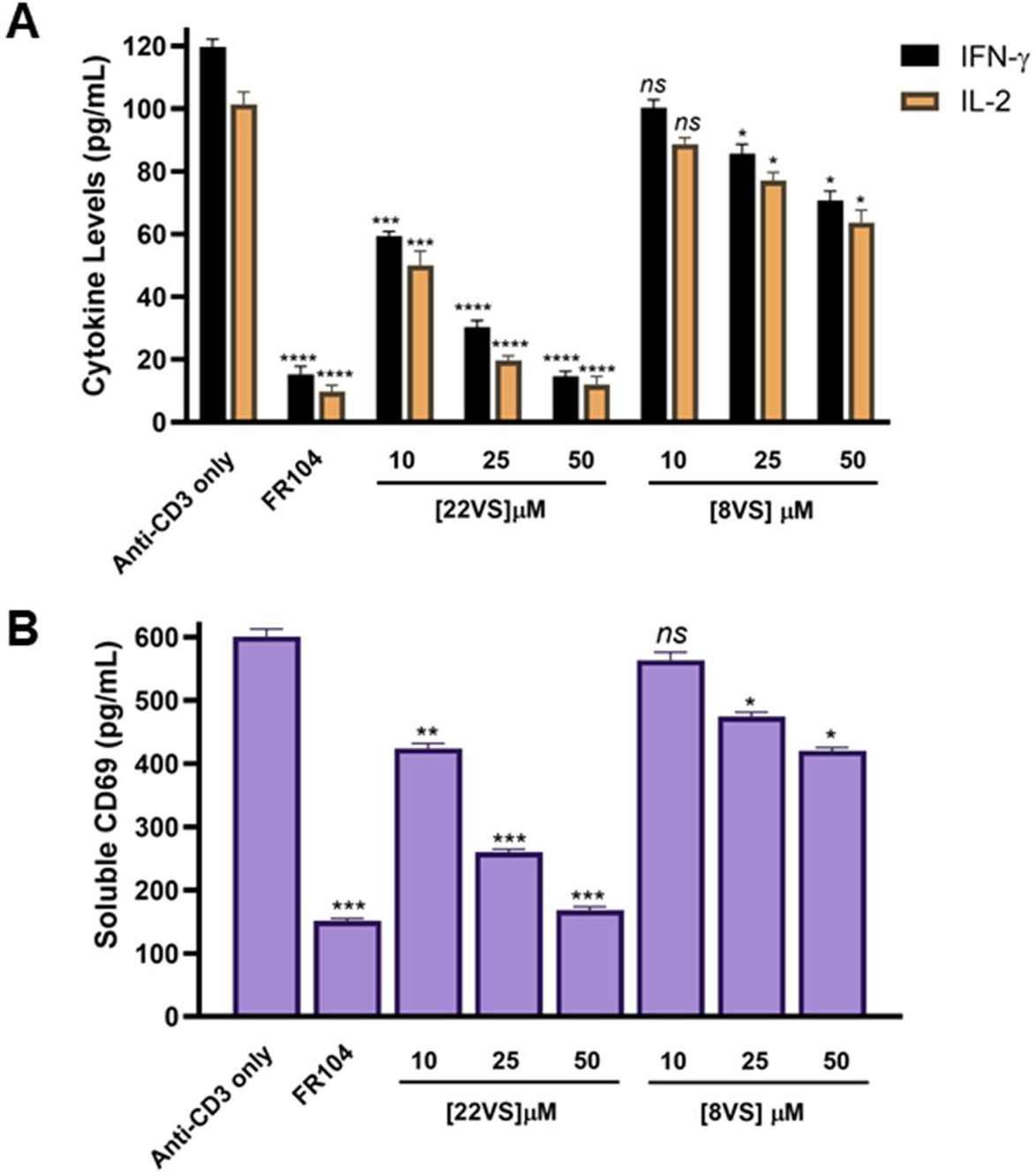
Dose-dependent inhibition of T cell activation markers in tumor–PBMC co-culture. **(A)** Soluble IFN-γ and IL-2 levels were quantified after 48-hour co-culture of A549 tumor spheroids with human PBMCs (E:T ratio 5:1) in the presence of anti-CD3 (0.3 µg/mL) and test compounds: FR104 (10 µg/mL), 22VS (10, 25, 50 µM), or 8VS (10, 25, 50 µM). **(B)** Soluble CD69 levels measured by ELISA after 48-hour co-culture of A549 tumor spheroids with human PBMCs (E:T ratio 5:1) in the presence of anti-CD3 (0.3 µg/mL) and test compounds: FR104 (10 µg/mL), 22VS (10, 25, 50 µM), or 8VS (10, 25, 50 µM). Data represent mean ± SD of n = 3 independent wells. Statistical comparisons to the “Anti-CD3 only” group were performed using one-way ANOVA followed by Dunnett’s post-hoc test. *ns* denotes non-significant, * *p* < 0.05, ** *p* < 0.01, *** *p* < 0.001, and **** *p* < 0.0001 relative to anti-CD3.

Compound 22VS demonstrated dose-dependent suppression of T cell activation. At 10, 25, and 50 μM, IFN-γ levels dropped to 60, 28, and 14 pg/mL, respectively; IL-2 to 50, 22, and 12 pg/mL; and sCD69 to 420, 260, and 160 pg/mL. At 50 μM, 22VS closely mirrored the FR104 profile, validating its potency and functional mimicry of a known antagonist (Figures 8A, 8B). In contrast, 8VS showed limited activity, with only modest reductions even at 50 μM (IFN-γ = 70 pg/mL, IL-2 = 62 pg/mL, sCD69 = 420 pg/mL), consistent with its weaker biochemical potency (IC50 = 60 μM).

Together, these findings establish the tumor–PBMC co-culture as a robust translational assay for assessing CD28-targeted modulation. The concordance between FR104 and 22VS supports the latter’s functional relevance, while 8VS’s minimal effect highlights the assay’s specificity. This system offers a scalable alternative to in vivo models for early immunomodulatory screening.

### 3.9 22VS Suppresses CD28-Dependent Cytokine Release in a Human PBMC–Mucosal Co-culture Model

We next assessed whether 22VS can suppress CD28-dependent immune activation in a physiologically relevant epithelial–immune interface. To this end, we employed a human PBMC–mucosal co-culture system using fully differentiated airway epithelial tissue (MucilAir™, Epithelix) overlaid with primary PBMCs and stimulated with anti-CD3 and soluble anti-CD28 (Figure 9A). Cytokine release into the apical compartment was measured after 48 hours.

**Figure 9.**
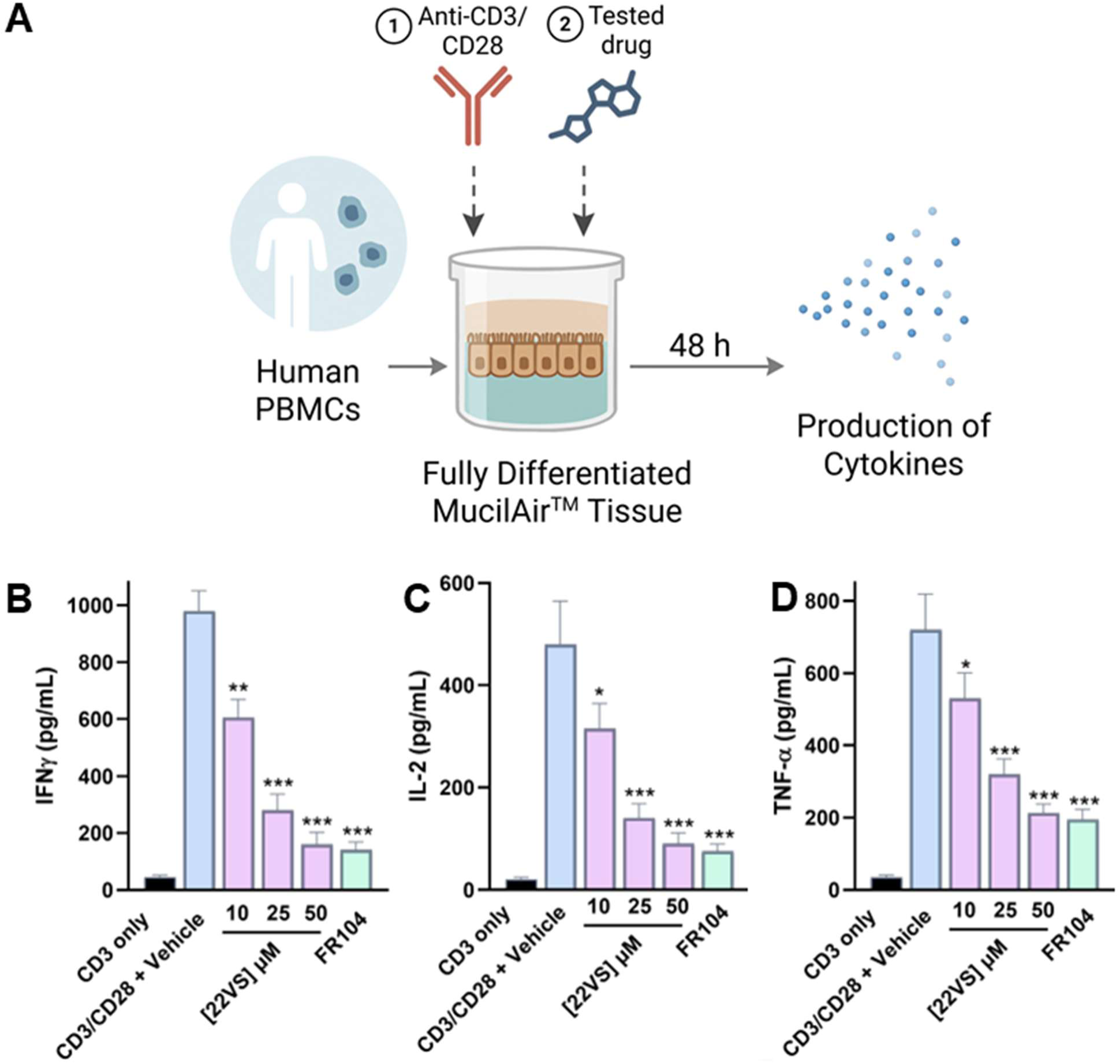
22VS suppresses CD28-dependent cytokine release in a human PBMC–mucosal co-culture model. **(A)** Schematic overview of the experimental setup. Human PBMCs were overlaid onto fully differentiated MucilAir™ tissue (Epithelix) and stimulated with plate-bound anti-CD3 and soluble anti-CD28 for 48 h in the presence of vehicle (0.1% DMSO), 22VS (10, 25, or 50 µM), or FR104 (10 µg/mL). **(B–D)** Quantification of IFN-γ, IL-2, and TNF-α secretion (pg/mL) in apical supernatants by ELISA. 22VS suppressed CD28-induced cytokine production in a dose-dependent manner, with levels at 50 µM comparable to FR104. Statistical comparisons to the “CD3/CD28 + Vehicle” group were performed using one-way ANOVA followed by Dunnett’s post-hoc test. * *p* < 0.05 and *** *p* < 0.001 relative to “CD3/CD28 + Vehicle” group.

Co-stimulation with anti-CD3/CD28 resulted in robust secretion of IFN-γ, IL-2, and TNF-α compared to anti- CD3 alone. Treatment with 22VS led to dose-dependent suppression of all three cytokines (Figure 9B). At 10 µM, 22VS reduced IFN-γ to 610 ± 55 pg/mL and IL-2 to 310 ± 40 pg/mL (Figure 9B). At 25 µM, the reductions were more pronounced (IFN-γ: 280 ± 40 pg/mL; IL-2: 140 ± 25 pg/mL), while at 50 µM, cytokine levels were near baseline (IFN-γ: 160 ± 30 pg/mL; IL-2: 90 ± 15 pg/mL). TNF-α showed a similar trend, decreasing from 720 ± 70 pg/mL in vehicle to 215 ± 20 pg/mL at the highest 22VS dose (Figure 9B). Notably, these effects were comparable to FR104 (10 µg/mL), which reduced IFN-γ, IL-2, and TNF-α to 140 ± 20, 75 ± 10, and 195 ± 18 pg/mL, respectively (Figure 9B). These results demonstrate that 22VS retains potent functional activity in a primary tissue model that recapitulates key features of human mucosal immunity, including epithelial–immune cross-talk, barrier integrity, and physiologically relevant cytokine gradients. The consistency of 22VS-mediated suppression across tumor–PBMC and mucosal co-culture models supports its mechanism as a bona fide CD28 pathway antagonist.

## 4. Discussion and Conclusion

Immune checkpoint and co-stimulatory receptors such as PD-1, CTLA-4, CD28, ICOS, and CD40 have emerged as critical modulators of T cell responses and are now established therapeutic targets in cancer and autoimmunity (Bates et al., 2023; Chen et al., 2025; Kennedy et al., 2024; Zhao et al., 2023). While biologics targeting these proteins—primarily monoclonal antibodies—have achieved clinical success, they carry inherent limitations, including high production costs, parenteral administration, potential immunogenicity, and restricted tissue penetration. These challenges have sparked growing interest in developing small-molecule modulators that can access intracellular and extracellular domains, offer oral bioavailability, and exhibit better tissue distribution and manufacturing scalability. Small molecule inhibitors of PD-L1 (e.g., BMS-1166 and CA-170), agonists of ICOS, and CD40–CD40L antagonists have all demonstrated encouraging results in preclinical and early clinical trials, affirming the feasibility of rationally designing drugs for immune receptors historically deemed “undruggable.”(Chen et al., 2020; Sasikumar et al., 2021). CD28, a homodimeric transmembrane receptor, is the canonical co-stimulatory molecule essential for full T cell activation. It engages B7 ligands (CD80 and CD86) on antigen-presenting cells to amplify TCR signals and promote IL-2 production, clonal expansion, and effector differentiation (Upadhyay, Kaur, et al., 2025). Although therapeutic antibodies like FR104 and lulizumab have shown promise in modulating CD28 signaling in models of autoimmunity and transplantation, small-molecule inhibitors remain poorly characterized. The primary challenge lies in the nature of the CD28–B7 interface: large, shallow, and dominated by polar contacts—features generally considered incompatible with classical drug-like compounds. However, recent progress in computational chemistry, fragment-based screening, and pocket detection algorithms has begun to reveal cryptic binding sites on similar immune proteins, suggesting that CD28 may also harbor exploitable topographies for small-molecule binding (Calvo-Barreiro & Gabr, 2025).

In this study, we employed a structure-guided virtual screening strategy to interrogate the surface of CD28 and identified a previously uncharacterized lipophilic canyon near residues L22, K24, Q25, P27, N129, G130 and I132. This pocket showed a propensity for binding ligands with both hydrophobic and hydrogen-bonding capabilities. Among the prioritized virtual hits, the 1,3,4-thiadiazole-containing scaffold stood out as a privileged chemotype, recurring across multiple docking poses and satisfying key pharmacophore features with a hydrogen bond interaction with the side chain of N129 being established by the 1,3,4-thiadiazole- containing scaffold. Based on these insights, we selected 33 hits for experimental validation and ultimately identified 22VS as the most promising lead compound. Biophysical characterization using TRIC and MST demonstrated that 22VS binds directly to CD28 with low micromolar affinity (Kd = 52.45 µM). This binding translated into functional inhibition of the CD28–CD80 interaction in ELISA assays (IC₅₀ = 7.80 µM). Notably, in a NanoBit-based cellular assay—which offers a more physiologically relevant model of membrane-bound protein–protein interactions—22VS disrupted both CD28–CD80 and CD28–CD86 engagement with submicromolar potency. The observed maximum inhibition was modest (∼30–40%), likely reflecting receptor saturation or limited target occupancy under cellular conditions. Nonetheless, the differential inhibition of CD86 over CD80 was consistent with their known differences in affinity toward CD28 and reinforces the selectivity of the compound (Collins et al., 2002; Kennedy et al., 2022). Importantly, 22VS exhibited no measurable cytotoxicity in Jurkat T cells up to 300 µM, distinguishing it from non-specific inhibitors and supporting its suitability for further development. Furthermore, in a CD28 Blockade Bioassay, 22VS inhibited T cell activation in a dose-dependent manner (IC₅₀ = 6.96 µM), indicating that its binding to CD28 functionally impairs downstream signaling. These data suggest that 22VS acts as a functional antagonist rather than a non-specific binder and supports the hypothesis that this previously unexploited binding pocket can be leveraged for immune modulation. Pharmacokinetic profiling supports 22VS as the lead candidate due to its superior drug-like properties. It demonstrated high solubility, favorable metabolic stability in both human and mouse liver microsomes, extended plasma half-life, and minimal CYP450 inhibition—together indicating a well-balanced ADME profile.

To assess immune modulation in a physiologically relevant context, we used a tumor–PBMC co-culture system mimicking key aspects of the tumor immune microenvironment, including TCR engagement and CD28-mediated co-stimulation. 22VS showed robust, dose-dependent suppression of T cell activation markers (IFN-γ, IL-2, and sCD69), closely matching the response elicited by the biologic CD28 antagonist FR104. In contrast, 8VS had minimal effect, reinforcing the importance of both biochemical potency and target engagement for effective immune modulation. These findings highlight 22VS as a promising lead for further preclinical development and demonstrate the utility of translational co-culture models for evaluating small-molecule immunotherapies targeting co-stimulatory pathways. The PBMC–mucosal co-culture results further support the translational potential of 22VS by demonstrating its activity in a primary human epithelial–immune interface. The compound showed dose-dependent suppression of CD28-driven cytokine release, closely mirroring the benchmark biologic FR104. These findings confirm that 22VS maintains efficacy in a physiologically relevant context that incorporates tissue-level features such as barrier function and immune–epithelial cross-talk. Importantly, this model extends the pharmacological validation of 22VS beyond suspension or tumor-based systems, highlighting its potential relevance for mucosal inflammatory diseases where CD28 costimulation is pathogenic. The consistency of inhibition across models underscores 22VS as a promising small-molecule CD28 antagonist with broad applicability.

Our findings are significant in the context of ongoing efforts to expand the pharmacological toolkit for modulating co-stimulatory pathways. Whereas PD-1/PD-L1 blockade has revolutionized cancer immunotherapy, a substantial proportion of patients fail to respond due to intrinsic resistance mechanisms, including compensatory upregulation of co-stimulatory signals like CD28 (Wang et al., 2023). Thus, selectively targeting CD28 may serve as an adjunct strategy to improve checkpoint inhibitor efficacy. In autoimmune and inflammatory disorders, CD28 antagonists may help suppress aberrant T cell activity without broadly impairing immune surveillance, offering a more targeted and tolerable approach than global immunosuppressants (Vanhove et al., 2017). The therapeutic relevance of CD28 inhibition is further underscored by parallels with other immune co-receptors. For example, small-molecule inhibitors of PD-L1 such as CA-170 have demonstrated oral activity and are currently in clinical trials. Similarly, peptide-based CD40 inhibitors are advancing through early-phase development (Solanki et al., 2023). These precedents validate the concept that even shallow or flexible interfaces can be modulated with appropriately designed molecules. Our study builds on this by identifying a novel binding site and delivering the first rationally designed, cell-active small molecule that inhibits CD28–B7 engagement with functional consequences in T cells. Despite promising results, several limitations remain. The current lead, 22VS, shows moderate potency and incomplete inhibition at saturating concentrations, likely due to solubility issues, suboptimal kinetics, or limited receptor accessibility. While docking and pharmacophore modeling support its binding mode, structural validation (e.g., crystallography or cryo-EM) is needed. Future efforts will optimize potency, solubility, and pharmacokinetics through iterative medicinal chemistry, informed by high-resolution structural data.

In conclusion, this study reveals a novel druggable pocket on the surface of human CD28 and introduces 22VS as a selective small molecule inhibitor that disrupts CD28–B7 interactions and suppresses downstream T cell activation. These findings provide compelling evidence that CD28 is amenable to pharmacological modulation via rationally designed ligands and establish a foundation for future efforts to optimize and develop CD28-targeted small-molecule therapeutics. As research continues to shift toward precision immunomodulation, our work underscores the value of integrating structure-based design with functional validation to unlock new targets and pathways in immune regulation.

## Abbreviations

PPI: Protein–Protein Interaction
MST: Microscale Thermophoresis
TRIC: Temperature-Related Intensity Change
ELISA: Enzyme-Linked Immunosorbent Assay
HTVS: High-Throughput Virtual Screening
SAR: Structure–Activity Relationship
IC50: Half Maximal Inhibitory Concentration
TCR: T Cell Receptor
MD Simulation: Molecular Dynamics Simulation

## Acknowledgments

This work was supported by the National Institute of Diabetes and Digestive and Kidney Diseases (NIDDK) under grant number R01DK137299 (PI: Gabr).

