## Supporting Information for "Small Molecule-Based Blockade of CD28 Suppresses T Cell Costimulation Across Cellular and Mucosal Co-culture Models"

### ^a^ Department of Radiology, Molecular Imaging Innovations Institute (MI3), Weill Cornell Medicine, New York, NY 10065, USA.

### ^b^ Freie Universität Berlin, Molecular Design Group, Institute of Pharmacy, Department of Biology, Chemistry & Pharmacy, Königin-Luisestr. 2+4, 14195 Berlin

**Contents**

| 1. | LCMS purity assessment of compound 22VS | **S2** |
| --- | --- | --- |
| **2.** | ^1^H NMR spectrum of compounds of 22VS | **S3** |
| **3.** | LCMS purity assessment of compound 8VS | **S4** |

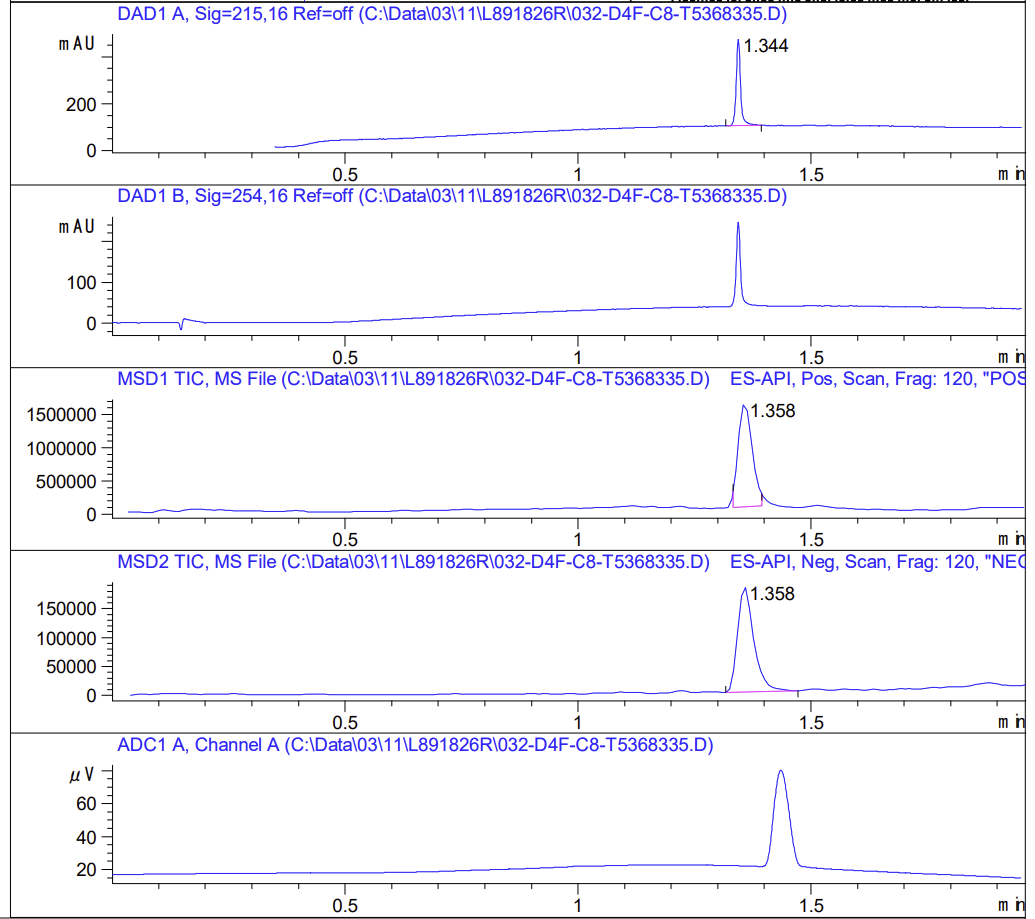

**Fig. S1.** LCMS purity assessment of compound 22VS.

**Fig. S2.** ^1^H NMR of 22VS.

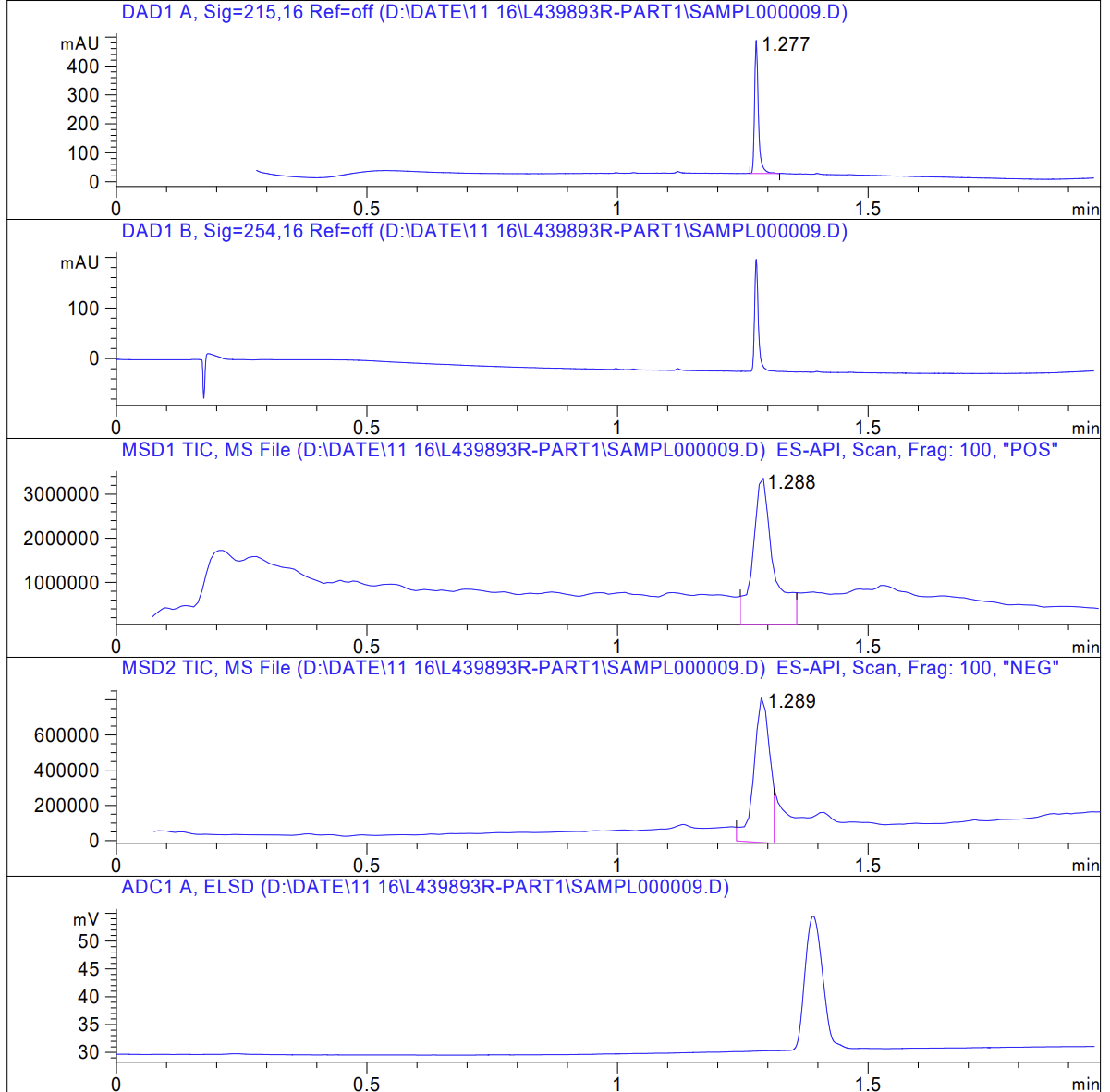

**Fig. S3.** LCMS purity assessment of compound 8VS.
